# Genomic surveillance and improved molecular typing of *Bordetella pertussis* using wgMLST

**DOI:** 10.1101/2020.10.28.360149

**Authors:** Michael R. Weigand, Yanhui Peng, Hannes Pouseele, Dane Kania, Katherine E. Bowden, Margaret M. Williams, M. Lucia Tondella

## Abstract

Multi-Locus Sequence Typing (MLST) provides allele-based characterization of bacterial pathogens in a standardized framework. However, current MLST schemes for *Bordetella pertussis*, the causative agent of whooping cough, seldom reveal diversity among the small number of gene targets and thereby fail to delineate population structure. To improve discriminatory power of allele-based molecular typing of *B. pertussis*, we have developed a whole-genome MLST (wgMLST) scheme from 214 reference-quality genome assemblies. Iterative refinement and allele curation resulted in a scheme of 3,506 coding sequences and covering 81.4% of the *B. pertussis* genome. This wgMLST scheme was further evaluated with data from a convenience sample of 2,389 *B. pertussis* isolates sequenced on Illumina instruments, including isolates from known outbreaks and epidemics previously characterized by existing molecular assays, as well as replicates collected from individual patients. wgMLST demonstrated concordance with whole-genome single nucleotide polymorphisms (SNP) profiles, accurately resolved outbreak and sporadic cases in a retrospective comparison, and clustered replicate isolates collected from individual patients during diagnostic confirmation. Additionally, a re-analysis of isolates from two statewide epidemics using wgMLST reconstructed the population structures of circulating strains with increased resolution, revealing new clusters of related cases. Comparison with an existing core-genome (cgMLST) scheme highlights the genomic stability of this bacterium and forms the initial foundation for necessary standardization. These results demonstrate the utility of wgMLST for improving *B. pertussis* characterization and genomic surveillance during the current pertussis disease resurgence.

## INTRODUCTION

Whooping cough (pertussis) is a respiratory disease with the highest rates of morbidity and mortality in young infants that continues to resurge in the United States (US) and many other countries. Vaccines against pertussis were introduced in the 1940s leading to dramatic reduction in reported disease incidence. However, the switch to acellular formulations in the 1990s was followed by increased reporting among all age groups in the decades since, despite high or increasing coverage with pertussis containing vaccines among industrialized countries (1). While not fully understood, resurgence likely results from multiple factors, including heightened awareness, expanded surveillance, improved diagnostics, shifting transmission dynamics, and pathogen evolution (1–4). Waning protection conferred by acellular vaccines is likely also responsible for increased disease among vaccinated individuals (5, 6).

Increased pertussis in the US manifests in local outbreak clusters, but also in cyclical, statewide and national epidemics.(7). Past molecular study of epidemics has been challenged by the low genetic diversity of *B.* pertussis, frequently described as a ‘monomorphic’ pathogen (8). Traditional multi-locus sequence typing (MLST) targeting select housekeeping genes provides very little discriminatory power and isolates of *B. pertussis* are often genotyped according to alleles of key vaccine immunogen-encoding genes, which resolve most isolates into only few sequence types (9–12). Quantifying specific repeat content with multilocus variable-number tandem-repeat analysis (MLVA) similarly reveals few discrete types (13). Alternatively, pulse-field gel electrophoresis (PFGE) has proven more useful for molecular typing, owing to the structural plasticity of the *B. pertussis* chromosome (14–16), but lacks throughput and standardization across laboratories (17, 18). The molecular study of poly-clonal epidemics, as well as broader geographic or temporal dynamics, has required a time-consuming combination of various methodologies as no single assay can sufficiently identify linked case clusters or describe the molecular characteristics of circulating *B. pertussis* to guide prevention and control efforts (9, 16, 19, 20).

Applications of high-throughput sequencing have transformed public health by exploiting the profound resolution of pathogen genomics for effective investigation and surveillance of infectious disease (21–24). A number of recent whole-genome sequencing (WGS) analyses of circulating *B. pertussis* have successfully reconstructed the accumulation of single nucleotide polymorphisms (SNPs) to reveal the bacterium’s population structure, geographic dispersion, and phylogenetic history with new depth (3, 25–29). However, allele-based molecular fingerprints may provide a more viable implementation of genome-based strain typing that can be standardized for routine use in public health laboratories (30–32). Whole-genome MLST (wgMLST) schemes, which capture the full complement of protein-coding genes in the genome, have been successfully applied to microbial pathogens for molecular epidemiology and food source attribution (22, 31, 33). More restrictive core-genome MLST (cgMLST) schemes, which evaluate only highly conserved genes, have also been implemented, including for *B. pertussis* (22, 34).

A growing number of recent *B. pertussis* clinical isolates recovered worldwide have been sequenced, including many closed assemblies. Here we leverage a collection of annotated, reference-quality genome assemblies to develop a standardized genome-based *B. pertussis* strain typing system using wgMLST and evaluate its performance with 2,389 sequenced isolates, primarily recovered from US pertussis cases. The curated scheme includes 3,506 protein-coding gene sequences, covering 81.4% of the average *B. pertussis* genome, and reproduced the population structure concordant with SNPs in retrospective analyses. These results highlight that the genomic stability of this bacterium perhaps makes wgMLST well suited for routine genome-based strain typing by public health institutions and pertussis researchers.

## RESULTS

### Locus curation

The wgMLST scheme was developed from the protein-coding genes predicted in closed, reference-quality genome assemblies from 214 *B. pertussis* isolates in the CDC collection combined with 11 publicly available genome sequences (TABLE S1). All multi-copy genes, paralogs, and IS-element transposases were excluded resulting in 3,681 orthologous loci captured in the initial version, the majority of which exhibited one allele. Each locus was evaluated with a larger set of raw sequencing reads to identify any systematic errors in allele calling due to either coding sequence (CDS) disruption (i.e., indels, gene truncations) or non-ACGT bases (i.e., Ns), as determined in BioNumerics. Locus reliability was further evaluated by confirming matching allele calls in sequencing reads from isolates which were independently confirmed to differ by <= 1 SNP using kSNP. The whole process of locus curation, described with detail in the methods, was repeated twice and removed 175 problematic loci. The final scheme included 3,506 loci, covering > 3.3 Mb (81.4%) of the *B. pertussis* genome and an average 76.8% of protein-coding nucleotides (FIGURE 1). Details of each locus are available in DATASET S1.

**Figure 1.**
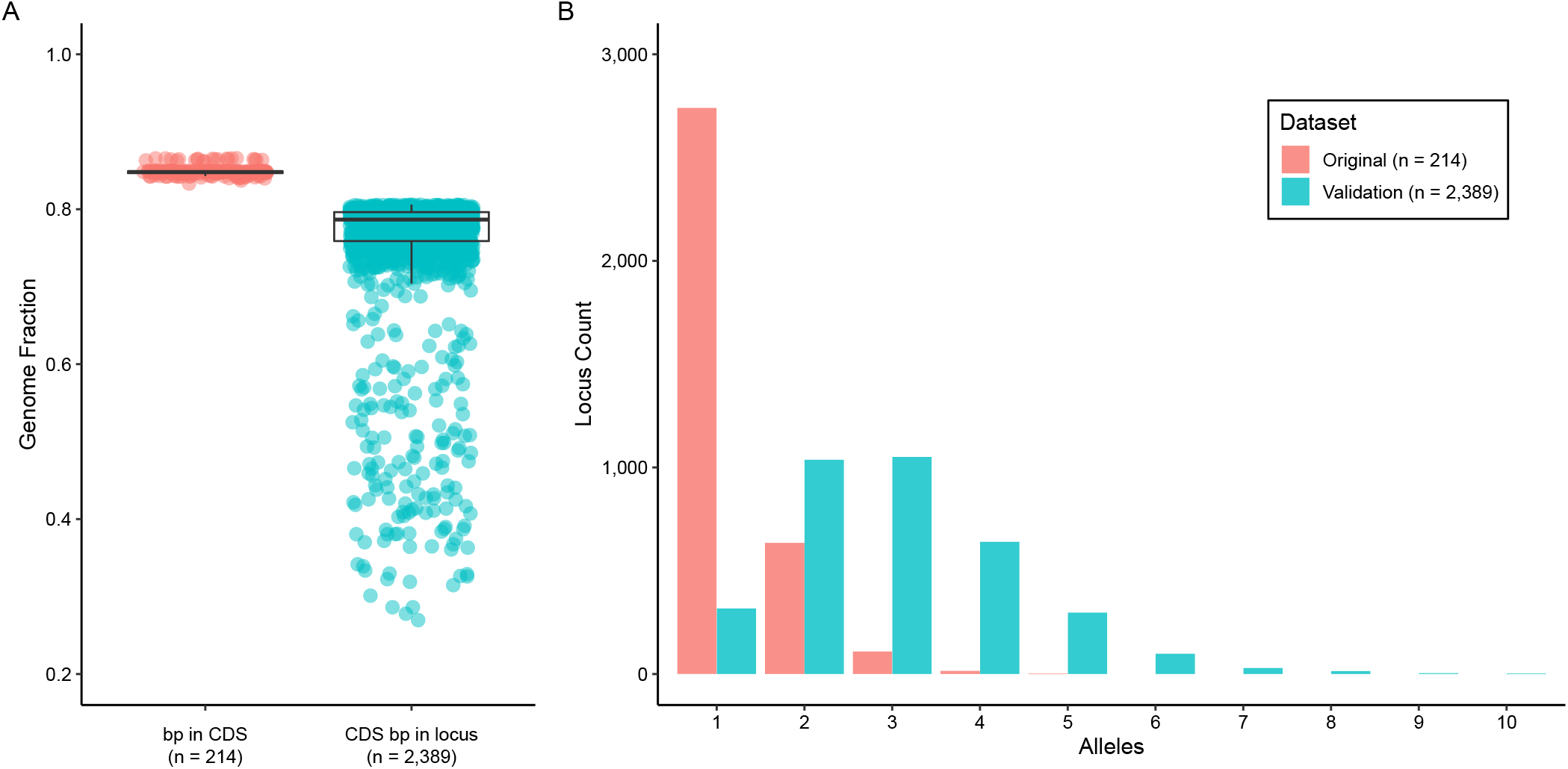
wgMLST scheme statistics. (A) The scheme was developed from 214 complete genome assemblies and covered an average 84.5% of each genome, while the final scheme captured an average 76.8% of protein-coding nucleotides in a large collection of sequenced isolates. (B) The distribution of unique alleles observed at each locus shifted as incorporating additional isolates revealed more diverse allele sequences at many gene loci.

### Performance testing

The process of allele calling in BioNumerics combines independent read mapping against a database (assembly-free, AF) and reference alignment to *de novo* assemblies (assembly-based, AB) to produce consensus allele calls. Performance was assessed across various metrics using sequencing reads from 2,389 isolates to determine potential variations in allele calling due to instruments, read lengths, coverage depth, or average read quality. Assembly quality proved a good indicator of allele calling, regardless of sequencing instrument or format, as better assemblies yielded more consensus allele calls (FIGURE S1). Read lengths influenced allele call performance more than sequencing instrument likely due to improved assembly as seen with 250 bp reads from either the MiSeq or HiSeq (FIGURE S1). Accordingly, decreased allele calling primarily resulted from non-ACGT errors that corresponded to the number of ambiguous bases in the contigs, further illustrating the dependence on *de novo* assembly (FIGURE S1, FIGURE S2). Failed allele calling due to CDS disruption did not depend on sequencing instrument or read length but rather reflected the known accumulation of pseudogenes present in *B. pertussis* genomes (35) (FIGURE S1, FIGURE S2). Locus-centric assessment of allele calling also indicated that the highest failure rates were due to CDS disruption in a small set of frequent pseudogenes while others were comparably sporadic. Based on these results, a minimum cutoff of 3,000 consensus allele calls was used for all subsequent analyses.

Allele profile differences among the 2,239 isolates with at least 3,000 allele calls were compared to pairwise SNPs distances predicted independently with kSNP to assess agreement between the two approaches for sequence-based strain typing. As expected, there was strong concordance between pairwise allele and pairwise SNP distances (FIGURE 2). Most sequenced isolates of *B. pertussis* differed by <100 SNPs but exhibited more allelic differences due to variations not linked to single base substitutions, which are not detectable with kSNP.

**Figure 2.**
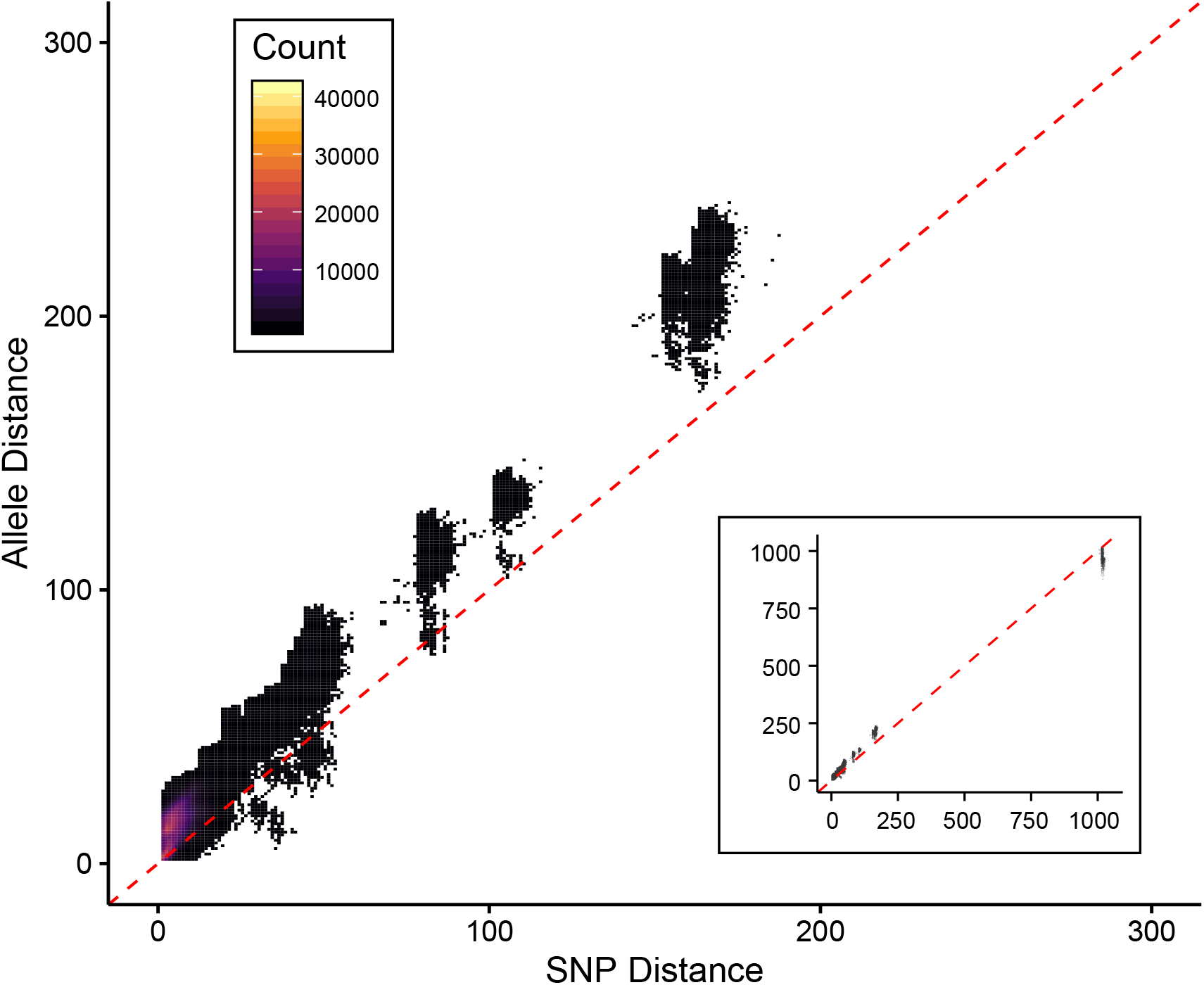
Concordance of pairwise distances. SNP and allele distances between all pairwise combinations of 2,389 sequenced isolates were concordant. Distances measured by wgMLST were consistently greater than measures of SNPs, reflecting the many types of sequence variation among the alleles. Distances are plotted as density according to the key and dotted lines indicate perfect correlation. Most isolates differed by < 200 alleles and a small number of very distant isolates resembled strain 18323 that differed from the majority by > 1000 alleles (inset).

### Reproducibility testing

To assess allele calling reproducibility and determine a minimum coverage depth cut-off, 10 sequencing read sets were randomly subsampled at seven coverage depths (9x, 14x, 21x, 31x, 46x, 70x, 105x), with five replicates. Each subsample was imported into BioNumerics and the total number of consensus allele calls, as well as their accuracy compared to the full read set, were compared across replicates. The total number of consensus allele calls remained above 3,000 for most replicates with average coverage depths >25x before dropping quickly (FIGURE 3A, FIGURE S4). As coverage decreased the consensus allele calls remained accurate, even as the total number of calls declined, and errors were only observed in two replicates at 9x depth (FIGURE 3B). Both errors were traced back to a single miscalled nucleotide in their respective loci. A minimum cut-off for average sequencing coverage depth was set at 30x for all subsequent analyses.

**Figure 3.**
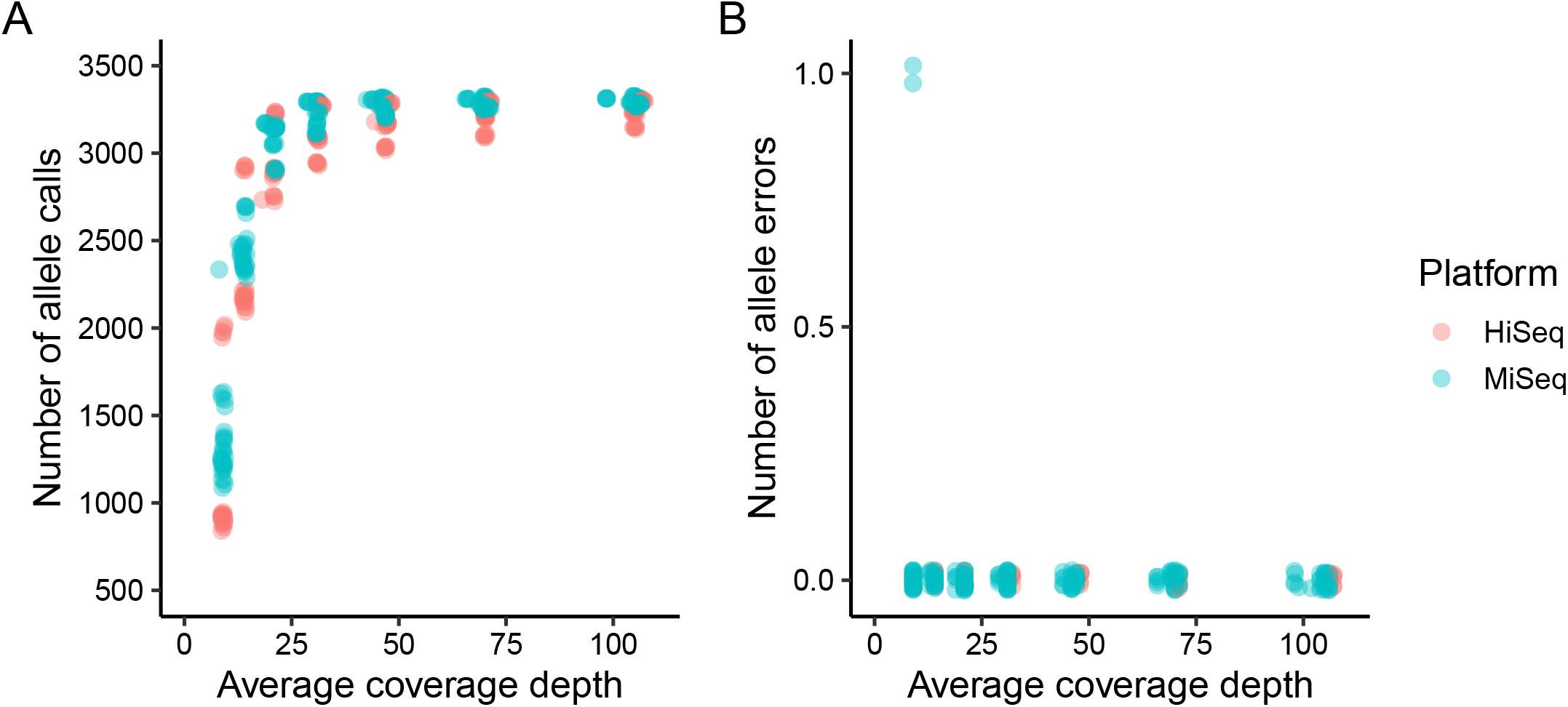
Technical replicates from subsampled read sets. At decreasing coverage depths (9x, 14x, 21x, 31x, 46x, 70x, 105x), replicated subsamples from select HiSeq and MiSeq read sets produced fewer consensus allele calls (A) but largely still made accurate allele calls compared to the full read set (B). Replicates for each read set are plotted separately in FIGURE S3.

Consistency among biological replicates was also evaluated using a collection of multiple isolates recovered from 152 individual patients. Select participating surveillance laboratories pick ‘sets’ of colonies (average = 5; range = 2-7) during culture confirmation and submission to CDC for characterization, including whole-genome sequencing (FIGURE S4). Sequence variation among isolates within each set was quantified as both SNPs and alleles. While 49 sets exhibited no differences by either measure, many sets include 1 SNP and pairwise distances up to 5 SNPs or 12 alleles were observed (FIGURE 4). Some allele differences resulted from mutations other than single base substitutions, such as indels.

**Figure 4.**
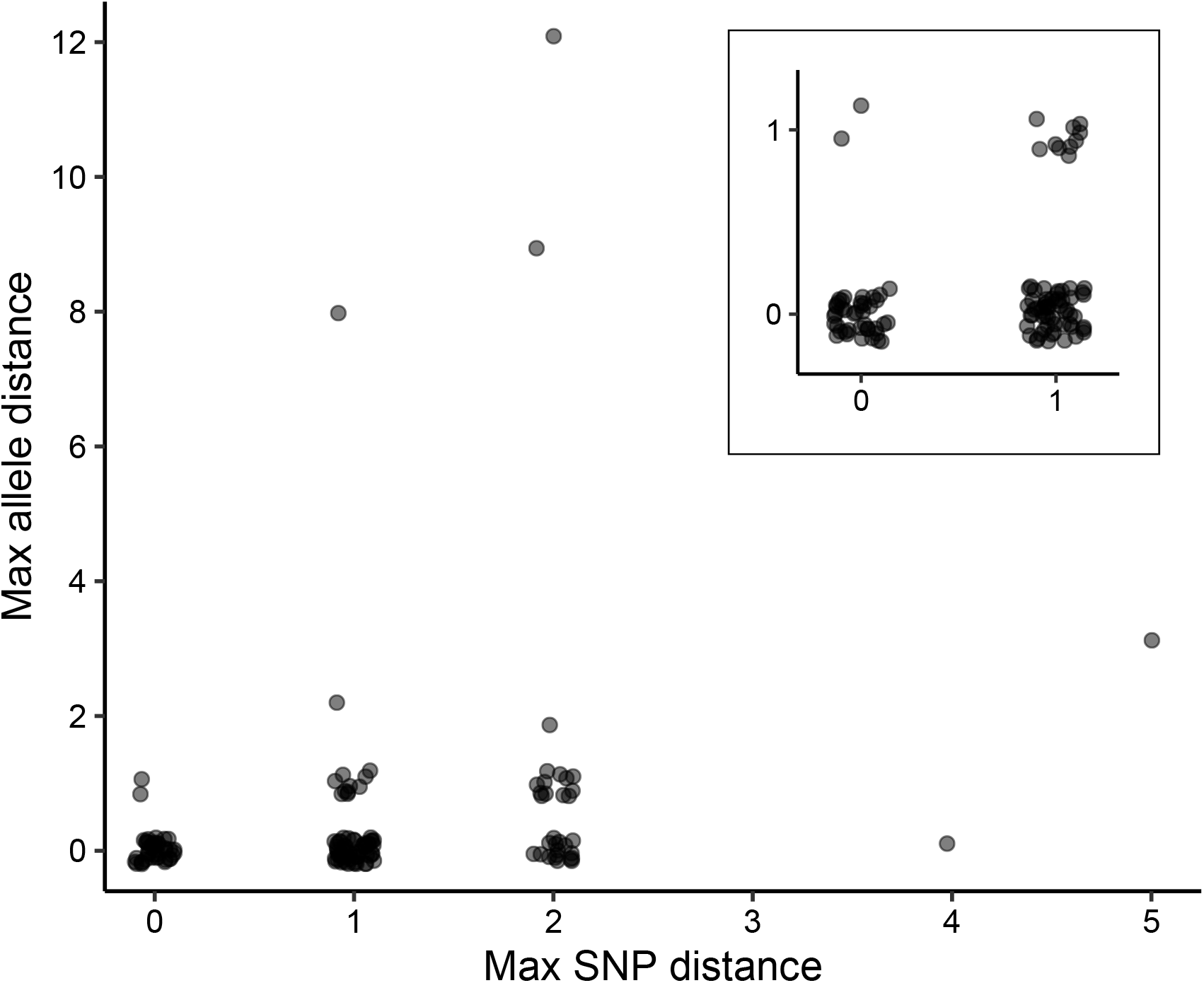
Biological replicates from individual patients. Sets of replicate isolates recovered from surveillance cases during culture confirmation were frequently not all identical, with a maximum pairwise distance within the set often equaling 1-2 SNPs or alleles. Inset shows sets with <= 1 SNP or allele maximum distance.

Taken together, these replicates indicate that allele calling results are reproducible and a minimum cut-off for sequencing depth was set at 30x for all subsequent analyses. Allelic and SNP variation detected within replicate isolate sets from individual patients suggests that genetic diversification occurs during infection. Therefore, the resolution of outbreak clustering may be limited to approximately 2 allele differences as most cases are represented by a single isolate and results could vary depending on which colony is selected during laboratory isolation.

### Retrospective outbreak cluster detection

To test the utility of the wgMLST scheme for studying the molecular epidemiology of pertussis, 12 isolates from epidemiologically linked cases associated with an outbreak occurring at a high school during a two-month period in 2016 were characterized. The case isolates were compared to 83 contemporaneous, sporadic isolates, which together represented 22% of cases reported from the surveillance catchment area that included the outbreak. The 12 outbreak cases shared an identical allele profile and were discretely clustered in a minimum spanning tree calculated from 157 variable loci (FIGURE 5). Comparing allele profiles also identified potential links to 3 case isolates recovered from infants, two of whom were siblings, which pre-dated the outbreak. All other contemporaneous isolates differed from the outbreak cluster by at least 2 alleles, some forming clusters of their own, demonstrating the effectiveness of wgMLST to delineate linked cases and potentially complement epidemiological investigation of localized *B. pertussis* outbreaks.

**Figure 5.**
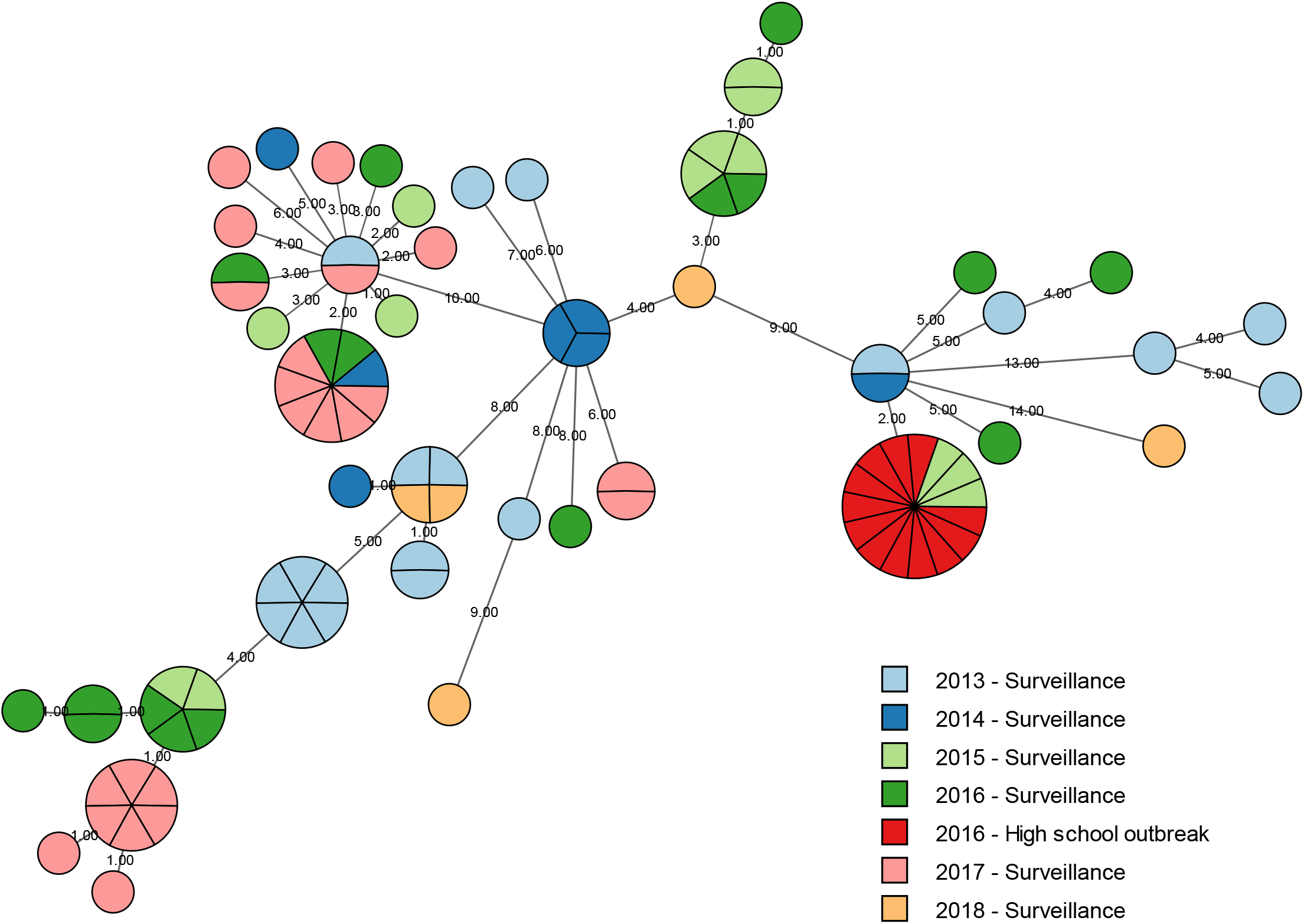
Molecular epidemiology of a high school outbreak. A minimum spanning tree calculated from 157 polymorphic loci clustered 12 case isolates from a high school outbreak, distinguishing them from 83 contemporaneous sporadic case isolates collected through routine surveillance in a retrospective comparison. Outbreak and sporadic case isolates are indicated according to the key. Node size indicates abundance and connecting lines are numbered according to allele distance.

### Population structure of statewide epidemics

Periods of increased disease have been reported across geographically defined regions (7), such as US states, and the test data here included sequenced *B. pertussis* isolates recovered from two such epidemics in Washington (9) and Vermont (36, 37). wgMLST revealed discrete population structures within each epidemic, confirming the polyclonal nature of each while identifying putative clusters of transmission among linked cases in a minimum spanning tree calculated from 186 variable loci (FIGURE 6). Some of the genotypes present during the epidemic were also detected among surveillance isolates collected after the epidemics. Combining the isolates from both state epidemics further confirmed that each included circulation of common genotypes, despite >3700 km of physical separation (FIGURE 6).

**Figure 6.**
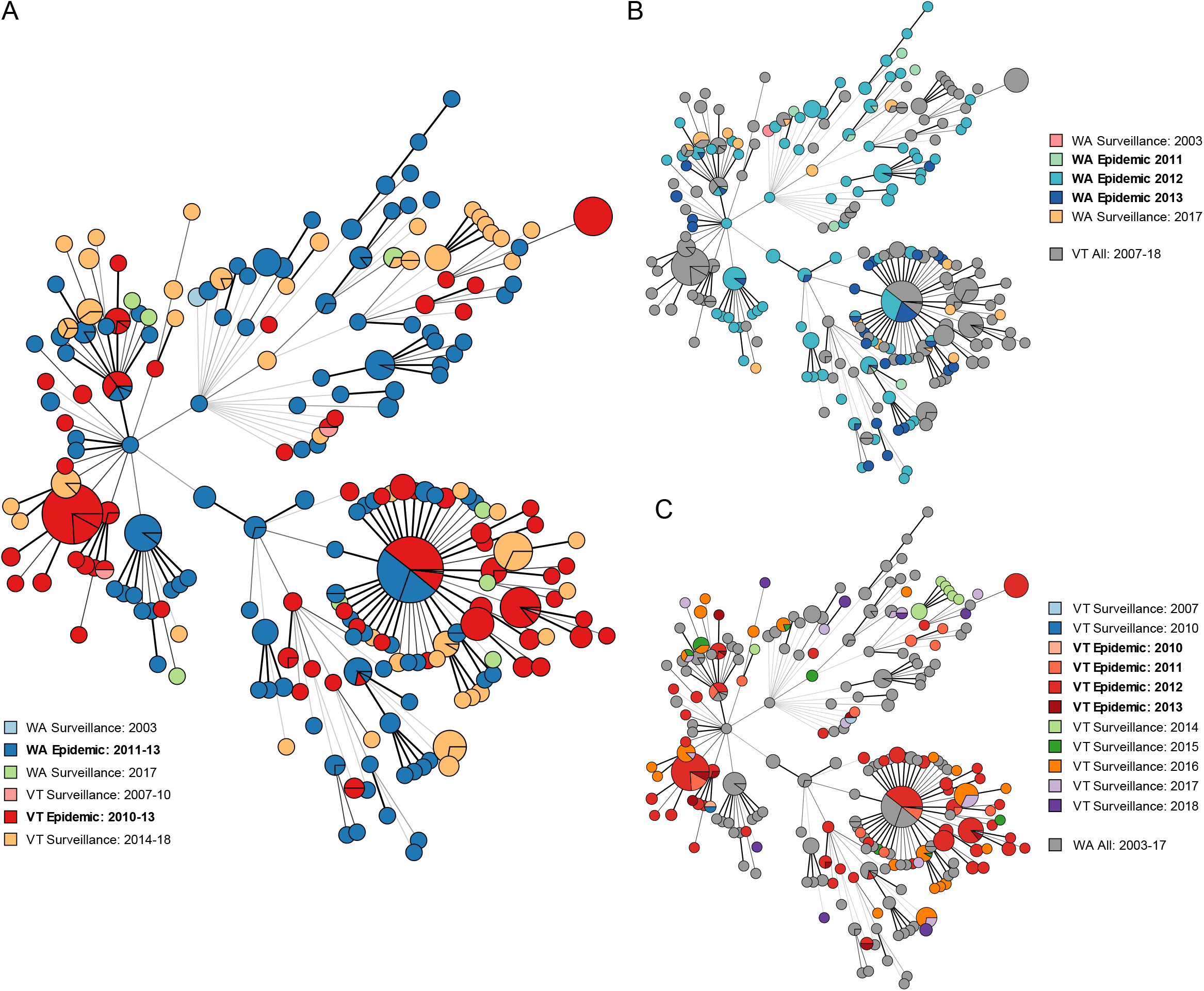
Molecular epidemiology of state-wide epidemics. Minimum spanning trees calculated from 186 core, polymorphic loci among 832 case isolates recovered from WA and VT. Many genotypes were common among the two epidemics (A) and both WA (B) and VT (C) likely included multiple clusters of transmission. Epidemic and routine surveillance case isolates are indicated according to the key in each panel. Node size indicates abundance and connecting lines are weighted according to allele distance.

### Comparison to Institute Pasteur cgMLST scheme

A similar core genome MLST (cgMLST) scheme for *B. pertussis* was recently developed at Institute Pasteur that includes 2,038 gene loci or 1.75 Mbp (42.7%) of the average *B. pertussis* genome (34). The overlapping gene content shared between that cgMLST scheme and the wgMLST scheme here was determined by reciprocal BLASTn alignment. The two schemes shared 1822 common loci, defined as >95% nucleotide sequence identity and >90% length overlap (FIGURE 7). Some loci in each scheme could not be directly linked, likely because the annotated genome inputs used for developing the two schemes relied on different gene prediction algorithms. Relaxing the minimum length overlap allowed matching an additional 108 shared loci. However, some predicted protein-coding gene loci in one scheme were split into two smaller genes in the other loci. After accounting for these gene prediction artefacts, there were 1583 unique wgMLST (45.2%) and 108 unique cgMLST (5.3%) loci that could not be matched (FIGURE 7). Thirty-three of these cgMLST loci did match predicted CDS in the input genomes here but were removed from the wgMLST scheme during curation. Many others aligned to predicted pseudogenes, all of which were excluded from the wgMLST scheme. Identified overlaps and unique gene loci are detailed in DATASET S1.

**Figure 7.**
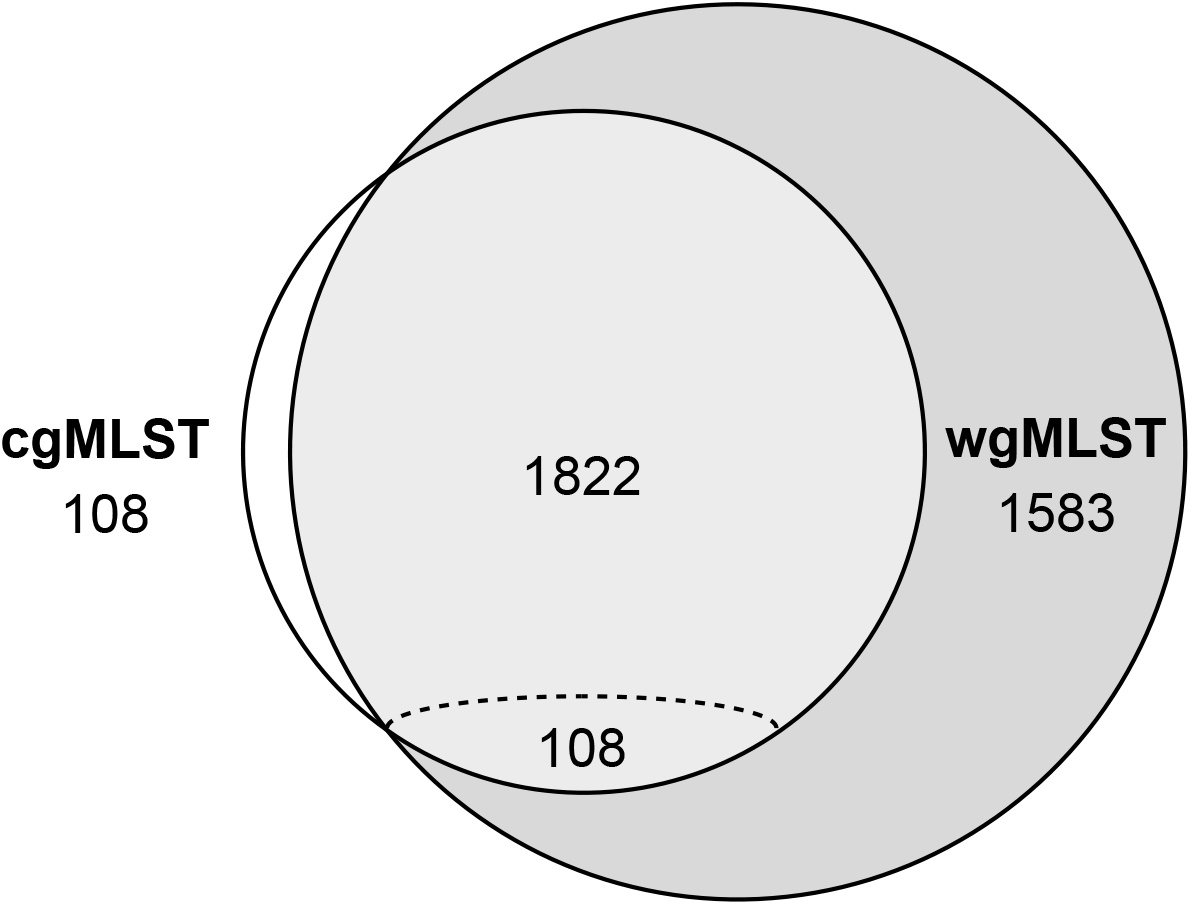
Gene content shared between the CDC wgMLST and Institute Pasteur cgMLST schemes. The two schemes shared 1822 orthologs and an additional 108 overlapping genes with varied lengths. Each scheme also included unique gene loci. Identified overlaps and unique gene loci are detailed in DATASET S1.

The resolution of cgMLST, implemented within BioNumerics to ensure consistent allele calling methodology, was tested using the same collection of high school outbreak and sporadic surveillance isolates above. A minimum spanning tree calculated from 76 polymorphic cgMLST loci (FIGURE 8) exhibited a similar topology to that determined using wgMLST (FIGURE 5) with subtle differences, as expected. However, cgMLST clustered the 12 outbreak isolates with an additional three surveillance isolates that differed by up to seven alleles according to wgMLST. The two schemes were further compared using pairwise allele distances among a subset of 379 sequenced isolates selected to represent the phylogenetic breadth of the larger collection. Similarly, the schemes were concordant but wgMLST identified more allelic differences among isolates reflecting the added resolution provided by the additional loci, as expected (FIGURE S5). The difference in resolution was particularly evident at shorter distances (Figure S5C) relevant for pertussis outbreak clusters delineation, consistent with observed clustering in the retrospective analysis (e.g. Figure 8).

**Figure 8.**
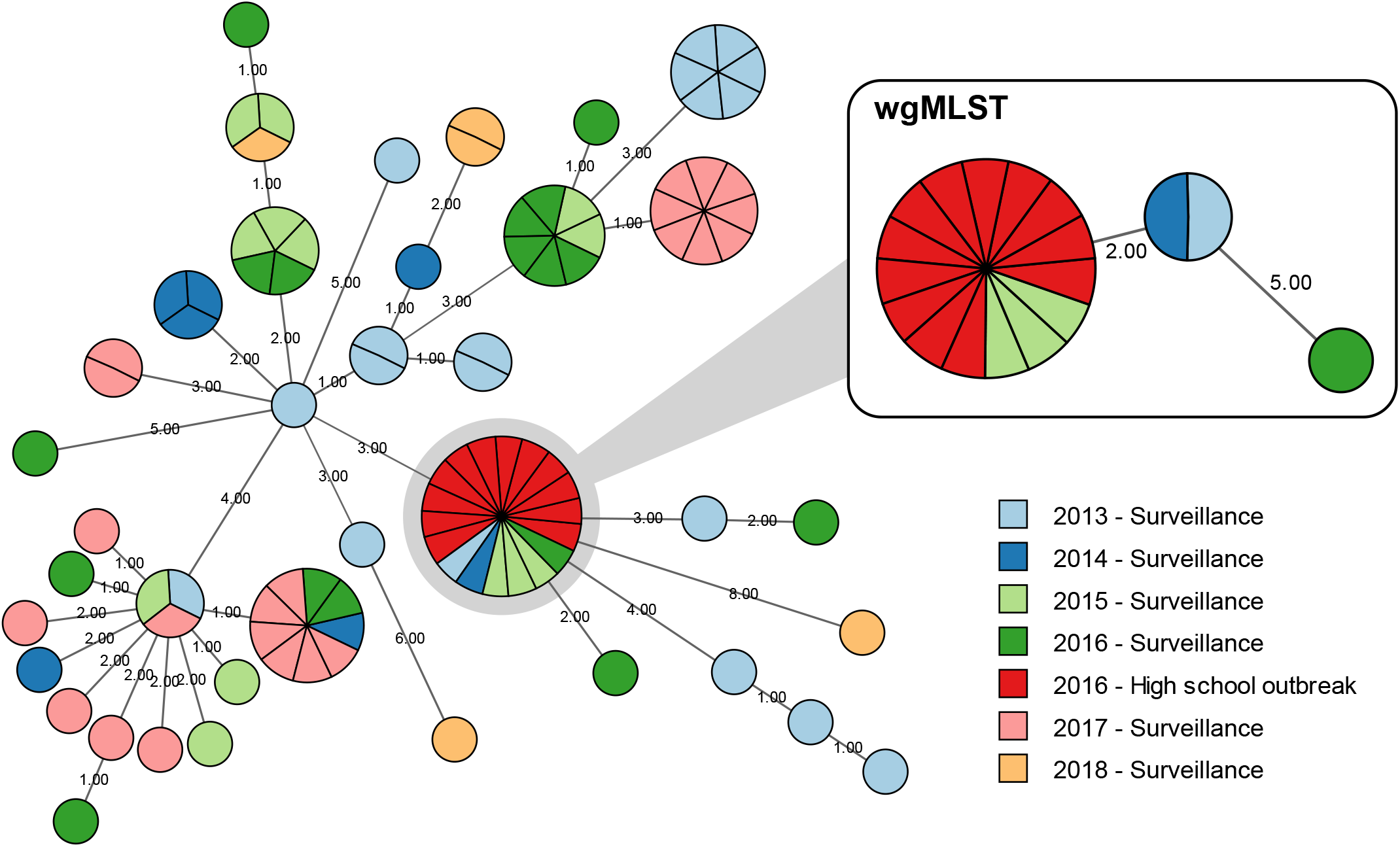
Typing resolution of cgMLST vs wgMLST. A minimum spanning tree from 76 polymorphic cgMLST loci clustered 12 case isolates from a high school outbreak with additional sporadic case isolates that were differentiated by wgMLST (inset). Outbreak and sporadic case isolates are indicated according to the key. Node size indicates abundance and connecting lines are numbered according to allele distance.

## DISCUSSION

In this study, we present the development and validation of a wgMLST scheme for *B. pertussis*, the primary agent of whooping cough. Traditional molecular methods provide little support for pertussis epidemiology and multiple assays used in combination have been needed to identify linkages among contemporaneous cases with only limited resolution (9, 12, 16). Through retrospective analyses, the results here demonstrate the utility of wgMLST for strain characterization using a single, genome-based assay within a standardized platform suitable for local and state public health laboratories. Widespread implementation of WGS and wgMLST for clinical *B. pertussis* can promote genomic surveillance, enhance understanding of the epidemiology of pertussis, and further empower study of pertussis resurgence.

The species *B. pertussis* has been frequently described as ‘monomorphic’ for its limited genome sequence diversity and nearly fixed accessory gene content (8). While such characteristics have made the traditional molecular characterization of clinical isolates challenging, the design approach of gene-by-gene allele typing may benefit from the large core fraction and lack of detectible recombination in the *B. pertussis* population. As a result, the wgMLST scheme developed here captures the majority of protein-coding nucleotides not associated with insertion sequence element (ISE) transposases, of which the *B. pertussis* genome harbors >250 (~7% of all CDS). Proliferation of these ISEs, particularly the >240 copies of IS*481*, facilitated genome reduction during the speciation of *B. pertussis* from the closely related ‘Classic Bordetellae’ (35, 38, 39). Such repetitive sequences also obstruct draft genome assembly from short-read sequencing data, which critically influences allele call performance by bioinformatic tools like BioNumerics when using read formats of 100 bp or less, regardless of average sequencing depth or read quality. Accordingly, appropriate wgMLST scheme curation considers the impact of both technical variables and microbe-specific biological idiosyncrasies, such as pseudogenes, repeat polymorphisms, and ISEs in the case of *B. pertussis*, as well as their intersection.

The economization of high-throughput sequencing and resolution of genome-based molecular typing has revolutionized characterization of numerous microbes associated with animal and human disease (21, 23). Successfully translating these technologies into application for molecular (genomic) epidemiology requires both standardization and portability. Perhaps the most successful example is the widespread implementation of wgMLST, supplanting PFGE, for foodborne pathogen surveillance (22, 40). Reported SNP phylogenetic reconstructions of *B. pertussis* clinical isolates have provided clear delineation of branching lineages and divergence from vaccine reference strains within the recent genomic history of *B. pertussis* (3, 15, 25). The data here highlight that wgMLST may provide additional resolution by capturing allele variants not resulting from single base substitutions and, therefore, provide a powerful single assay for strain typing and molecular epidemiology of *B. pertussis*.

A similar cgMLST scheme for *B. pertussis* was recently developed and reported by Institut Pasteur (34). That scheme targets genes present in nearly all isolates (‘core’) in contrast to the larger wgMLST scheme here. Comparing the two schemes revealed that they differed beyond the number of loci and the cgMLST scheme was not simply a subset of wgMLST. Differences in input data and gene prediction algorithms used to develop the two schemes produced CDS discrepancies, highlighting subtle differences in popular gene finding approaches (41, 42). Accurate locus detection and subsequent allele calling, not just in BioNumerics, benefits from conserved start and stop codon positions (31). The comparison here suggests that wgMLST does provide additional resolution in pairwise measurements, particularly among closely related *B. pertussis* isolates separated by distances relevant for outbreak cluster delineation. Broad application of allele-based typing for molecular epidemiology and genomic surveillance would benefit from scheme harmonization, which will require careful modification of loci in both schemes, starting with those which overlap only under relaxed alignment parameters. Such efforts would surely be rewarded with a more thorough database of observed, circulating allelic variation for use by varied public health institutions and researchers, including those focused on developing future pertussis vaccines.

An allele-based approach to strain characterization cannot resolve chromosome structure variation, which provides a significant source of genomic diversity among circulating *B. pertussis* (14, 15, 43). Genomes of *B. pertussis* clinical isolates exhibit frequent rearrangement, most often as large inversions, but more recently also amplifications (28, 44). It remains unclear whether such structural forms of genomic variation yield phenotypes, such as varied transmission or clinical disease presentation, but observed patterns among circulating isolates compared to common reference strains suggests they are under selection (45). These types of genomic structural features remain largely intractable, particularly by short-read sequencing platforms widespread in public health settings. However, the example retrospective datasets presented here, and previously (34), demonstrate the utility of allele-based strain typing for linking cases based on inferred ancestral relationships among recovered *B. pertussis* isolates. Previous comparative study of rearrangement variation among closed genome assemblies has revealed that many chromosome structures are phylogenetically restricted (15, 45), suggesting that reconstructing *B. pertussis* populations from polymorphic SNPs, or alleles, still captures meaningful relationships among case isolates.

Perhaps the largest barrier to widespread implementation of wgMLST (or cgMLST) for genomic surveillance of *B. pertussis* is the continued decline of diagnostic culture. All the data included here were derived from whole-genome sequencing of cultured isolates but on average fewer than 3.5% of US cases captured annually by the Enhanced Pertussis Surveillance/Emerging Infections Program (EPS) yield isolates (46). In principle, wgMLST can be applied to data derived from direct – ‘metagenomic’ – sequencing of clinical nasopharyngeal specimens, but will likely require careful modification. Limited observation of allelic variation among replicate isolates here highlight that application of wgMLST to direct sequencing data will need to evaluate polymorphic loci. For example, at least some replicate isolates recovered from individual patients differed by more alleles than were used to delineate a retrospective high school outbreak. Defining cluster cut-offs for sequence-based typing is a common problem (22, 30, 32), made more challenging by within-patient sequence variability of an organism with so little diversity. Solving these challenges and successful interoperability of wgMLST and direct sequencing is likely the only way for this, or any other method of genomic surveillance, to advance the study of pertussis resurgence. Hopefully the result facilitates production of sufficient datasets to enable large-scale, integrated analysis of genomic and epidemiological data.

## MATERIALS AND METHODS

### Strain selection

The Centers for Disease Control and Prevention’s (CDC) collection includes US *B. pertussis* isolates gathered through routine surveillance and during outbreaks. In total, sequence data from a convenience sample of 2,389 isolate genomes were included in the current study based on availability and most were selected for sequencing as part of previous studies (Table S2). Many isolates were obtained through the Enhanced Pertussis Surveillance/Emerging Infection Program Network (EPS) (46), including sets of 2-7 replicate isolates (average = 5) recovered from 153 patients during diagnostic culture confirmation.

### Genomic DNA preparation and sequencing

Isolates were cultured on Regan-Lowe agar without cephalexin for 72 h at 37 C. Genomic DNA isolation and purification was performed using the Gentra Puregene Yeast/Bacteria Kit (Qiagen; Valencia, CA) with slight modification (36). Briefly, two aliquots of approximately 1 × 10^9^ bacterial cells were harvested and resuspended in 500 uL of 0.85% sterile saline and then pelleted by centrifugation for 1 min at 16,000 × g. Recovered genomic DNA was resuspended in 100 uL of DNA Hydration Solution. Aliquots were quantified using a Nanodrop 2000 (Thermo Fisher Scientific Inc.; Wilmington, DE). Whole genome shotgun libraries were prepared using the NEB Ultra Library Prep kit (New England Biolabs; Ipswich, MA) for sequencing on either the MiSeq or HiSeq (Illumina; San Diego, CA). Sequencing reads from 10 isolates were randomly selected and subsampled without replacement yielding 5 replicate samples at each of 7 coverage depths (9x, 14x, 21x, 31x, 46x, 70x, 105x).

### wgMLST scheme design

The initial wgMLST scheme was developed from all protein-coding genes predicted in closed, reference-quality genome assemblies from 214 *B. pertussis* isolates (TABLE S1). Multi-copy genes and paralogs (e.g. ISE transposases) with > 95% sequence identity were detected and removed, excluding 240-260 CDS (6.5%) per genome. The remaining CDS were clustered into 3,681 orthologous loci. Each locus was further evaluated based on consensus allele call frequency and errors using custom scripts from Applied Maths, as well as manual inspection of allele alignments in BioNumerics. Loci were manually removed from the scheme based on criteria such as low frequency, low-complexity sequence repeats, homopolymeric tracts, length discrepancy, or variable allele calling between replicates. The process of locus curation was repeated twice, first with initial input set of 214 genomes and then with a larger collection of 614 isolates, leaving 3,506 loci in the final scheme.

### wgMLST allele calling and strain comparison

Allele calling was performed with the BioNumerics (v7.6.3) Calculation Engine. Imported sequencing reads were quality trimmed and filtered (min average read quality = 25, min read tail quality score = 15, min read length = 35 bp) before *de novo* assembly using Spades v3.7.1 (careful mode, min contig length = 300 bp) (47). Consensus allele calls were derived from the combination of read mapping (assembly-free, AF; k = 35; min coverage = 3x, min forward = 1x, min reverse = 1x) and reference alignment to the assembled contigs with discontinuous MegaBLAST (assembly-based, AB; k = 11, min similarity = 95%, allow gapped alignment, start/stop codon hunting off). Allele pattern comparisons were performed by selecting all sequencing read sets with at least 3,000 consensus allele calls and filtering out any monomorphic loci. Pairwise distances were determined using a simple cluster analysis based on categorical differences and UPGMA hierarchical clustering in BioNumerics. Minimum-spanning trees were calculated in BioNumerics using the advanced cluster analysis for categorical data.

### SNP detection

SNP variation among sequenced isolate genomes was determined with the exported *de novo* assemblies from BioNumerics using kSNP3 with k = 23 (48). Pairwise distances were calculated from all variable SNPs shared between each pair of sequenced isolates.

### Comparison to Institute Pasteur’s cgMLST scheme

The cgMLST scheme and allele definitions developed at Institute Pasteur (34) were kindly provided by Sylvain Brisse and Valérie Bouchez. Ortholog matching between the cgMLST and wgMLST schemes was performed by reciprocal best-match alignment using BLASTn (minimum 95% identity and 90% length match). Unmatched loci were further compared to identify overlapping sequence content with relaxed alignment parameters (minimum 90% identity and 50% length match).

The cgMLST scheme was loaded into a local database in BioNumerics and allele calling was performed with select isolates using the same parameters as indicated above for wgMLST. A representative subset of sequenced isolates was derived from the collection of 2,039 read sets with at least 3,000 wgMLST consensus allele calls by clustering isolates with 0-1 pairwise SNPs using mcl (49). One isolate was selected from each of the resulting clusters and combined with all unclustered (unique) isolates, as well as isolates from a retrospective high school outbreak and replicates from individual patients, into a dataset of 379 isolates representing the phylogenetic breath of the larger collection.

## Data availability

The whole-genome shotgun sequences are available from the NCBI Sequence Read Archive, organized under BioProject accession number PRJNA279196.

## Acknowledgements

We thank Pam Cassiday and Tami Skoff (CDC), Lingzi Xiaoli and Matt Cole (IHRC, Inc.), the CDC Biotechnology Core Facilities Branch Genome Sequencing Laboratory, The Enhanced Pertussis Surveillance/Emerging Infection Program Network, and Sylvain Brisse and Valérie Bouchez (Institut Pasteur).

This work was made possible through support from CDC’s Advanced Molecular Detection (AMD) program.

The findings and conclusions in this report are those of the authors and do not necessarily represent the official position of the Centers for Disease Control and Prevention. Use of trade names and commercial sources is for identification only and does not imply endorsement by the Centers for Disease Control and Prevention, the Public Health Service, or the U.S. Department of Health and Human Services.

